# Human stem cell transplantation for Parkinson’s disease: A systematic review of *in situ* survival and maturation of progenitors derived from human embryonic or induced stem cells in Parkinsonian models

**DOI:** 10.1101/2024.03.28.587203

**Authors:** Giulia Comini, Eilís Dowd

**Affiliations:** Pharmacology & Therapeutics and Galway Neuroscience Centre, University of Galway, Ireland

**Author notes:** **Corresponding author:** Professor Eilís Dowd.

**Keywords:** Parkinson’s disease, Cell replacement therapy, Embryonic stem cells (ESCs), Induced pluripotent stem cells (iPSCs), Transplant, Graft, Survival, Differentiation

## Abstract

Stem cell-based brain repair is a promising emergent therapy for Parkinson’s which is based on years of foundational research using human fetal donors as a cell source. Unlike current therapeutic options for patients, this approach has the potential to provide long-term stem cell-derived reconstruction and restoration of the dopaminergic input to denervated regions of the brain allowing for restoration of certain functions to patients. The ultimate clinical success of stem cell-derived brain repair will depend on both the safety and efficacy of the approach, and the latter is dependent on the ability of the transplanted cells to survive and differentiate into functional dopaminergic neurons in the Parkinsonian brain. Because the pre-clinical literature suggests that there is a considerable variability in survival and differentiation between studies, the aim of this systematic review was to assess these parameters in human stem-derived dopaminergic progenitor transplant studies in animal models of Parkinson’s. To do so, a defined systematic search of the PubMed database was completed to identify relevant studies published up to March 2024. After screening, 76 articles were included in the analysis from which 178 separate transplant studies were identified. From these, graft survival could be assessed in 52 studies and differentiation in 129 studies. Overall, we found that graft survival ranged from <1% to 500% of cells transplanted, with a median of 51% of transplanted cells surviving in the brain; while dopaminergic differentiation of the cells ranged from 0% to 46% of cells transplanted with a median of 3%. This systematic review suggests that there is considerable scope for improvement in the differentiation of stem cell-derived dopaminergic progenitors in order to maximize the therapeutic potential of this approach for patients.

## Introduction

The concept of human stem cell-derived brain repair for Parkinson’s disease is built on decades of research using fetal cells as a source of healthy cells for transplantation [1]. The first studies in this field date back to the 1970s when groups in the US [2] and Sweden [3] pioneered this approach using rat-to-rat fetal transplants. These studies demonstrated that fetal dopamine neuron-rich tissue transplanted into the brain could survive and ameliorate motor dysfunction in dopamine-depleted rats. A decade or so of preclinical research followed leading to the first clinical trials of human fetal transplantation in patients with Parkinson’s disease [4, 5].

Over the intervening decades, numerous preclinical studies and clinical trials provided proof-of-principle for the efficacy and safety of cellular brain repair, with fetal dopaminergic grafts demonstrating the ability to survive in the brain, reinnervate the dopamine-depleted striatum, and ameliorate dysfunction/symptoms in Parkinsonian models and patients [reviewed in 1, 6]. However, it also became apparent that this approach had serious limitations associated with the need for human fetal donors including supply, ethical concerns and logistical difficulties, all of which would ultimately preclude this translating to a therapy for patients.

In parallel with these efforts, embryonic stem cells (ECSs) were first cultured from mice [7, 8] and later humans [9], and protocols to convert these to a dopaminergic phenotype were later developed and refined [reviewed in 10]. This led to the first mouse-to-rat [11] and human-to-rat [12] studies of ESC-derived brain repair in Parkinsonian rats. With the discovery that somatic cells could be reprogrammed into induced pluripotent stem cells (iPSCs; [13–15], the era of stem cell-derived brain repair for Parkinson’s disease was truly underway [16, 17]. In contrast to fetal transplantation, both ESCs and iPSCs offered the possibility of a limitless supply of quality-controlled dopaminergic neurons for allogenic (donor-to-patient) transplantation, with the iPSCs offering the additional potential of autologous (patient-to-self) transplantation.

Over the last 20 years, numerous preclinical trials have been undertaken to determine the efficacy and safety of human stem cell-derived brain repair in Parkinsonian models, and the outcome of these has led to the initiation of 4 clinical trials that are currently underway (Kyoto Trial: UMIN000033564; BlueRock Trial: NCT04802733; STEM-PD: NCT05635409; S.Biomedics Trial: NCT05887466). Critically, the ultimate success of these and future clinical trials is highly dependent on the ability of the transplanted stem cell-derived dopaminergic progenitors to survive and differentiate into functional dopaminergic neurons in the Parkinsonian brain. Although *in situ* survival and differentiation of ESC and iPSC-derived dopaminergic progenitors is well established in rodent and primate models of Parkinson’s disease, this is highly variable between, and even within, studies.

Therefore, because of the pivotal role that *in situ* survival and dopaminergic differentiation will play in the clinical success of stem cell-derived brain repair for Parkinson’s, the aim of this review was to systematically assess these parameters in human stem-derived dopaminergic progenitor transplant studies in animal models.

## Methods

### Search Strategy

This study was completed in accordance with the PRISMA 2020 guidelines [18] and used a search strategy in the “PUBMED” database to find articles related to human stem-derived dopaminergic progenitor transplantation in animal models of Parkinson’s. The specific search string used was (stem AND (embryonic OR induced) AND Parkinson’s AND dopaminergic AND (transplant or graft)). This search identified a total of 887 articles spanning 1997 to March 2024. These articles were then screened according to the search strategy outlined below and depicted in the PRISMA flow diagram (Fig. 1). This yielded 76 articles which were included in this systematic review. All the articles obtained were managed using Microsoft Excel and EndNote software.

**Fig. 1.**
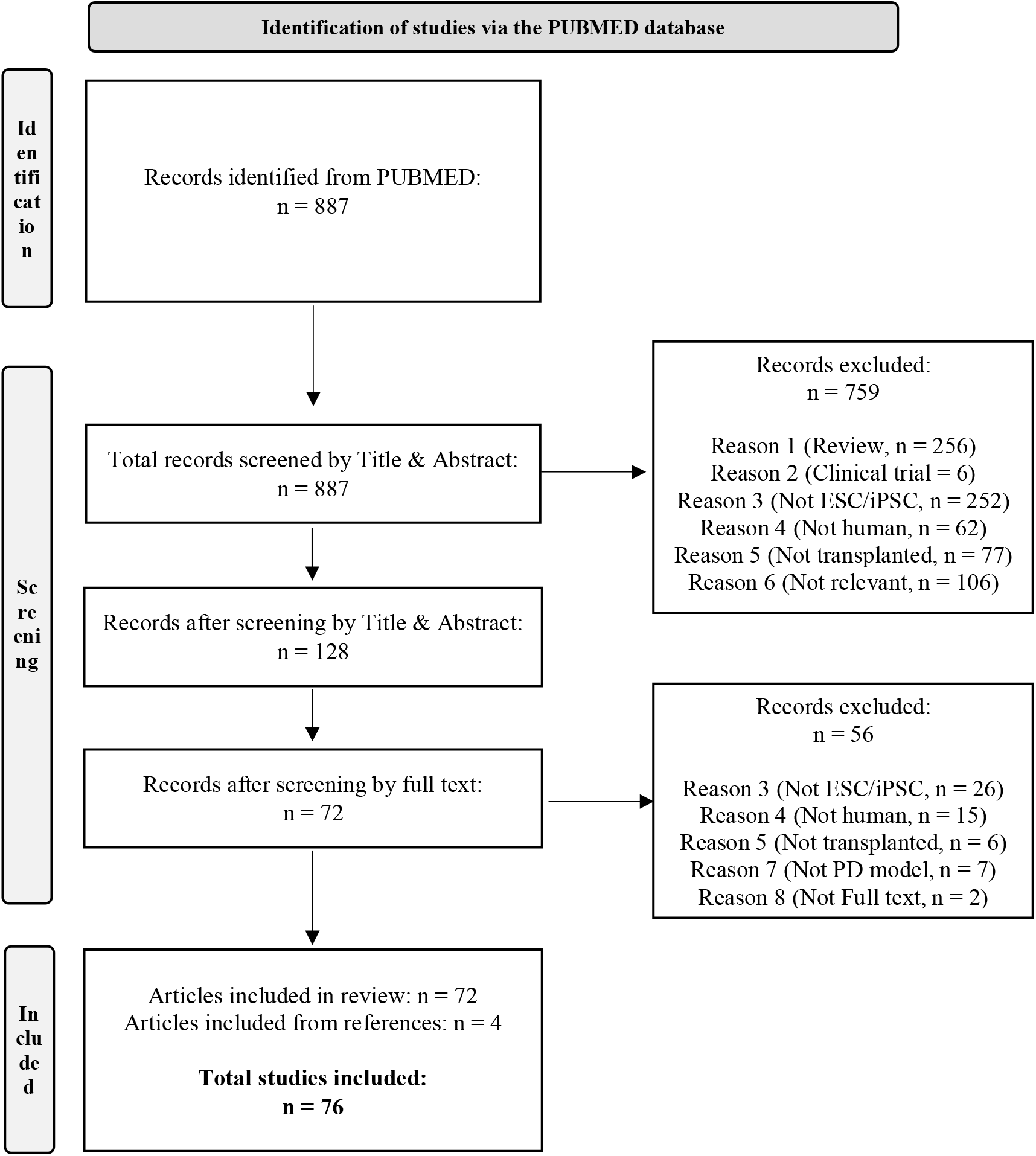
PRISMA flow chart. Flow chart based on PRISMA 2020 guidelines [18] detailing the screening strategy employed for the study selection in the present study.

### Screening Strategy

The 887 articles identified were first screened by Title and Abstract and the following inclusion and exclusion criteria were applied. Inclusion criteria: 1) original research studies, 2) human ESC or iPSC-derived dopaminergic progenitors, 3) transplanted into the brain. Exclusion criteria: 1) review articles, 2) clinical trials, 3) not ESC or iPSC cells, 4) not human origin, 5) no transplantation, 6) not relevant for other miscellaneous reasons. This initial screen left 128 articles which were screened by full text. In this second screen, two additional exclusion criteria were used: 7) transplanted animals were not Parkinsonian, and 8) full text was not available. This left 72 articles for inclusion to which a further 4 were added that were identified from the cited publications in the selected articles.

### Data Extraction

Variables manually extracted from the selected articles (Table 1 and references therein) included stem cell type (ECSs or iPSCs), cell line, day of differentiation at time of transplantation (for protocols using dual SMAD inhibition only), host species, Parkinsonian model, immunological status of the host (i.e. immunosuppressed vs. immunodeficient), week of sacrifice after transplantation, number of cells transplanted, number of cells surviving and number of cells differentiating into dopaminergic neurons. For cell survival, cell counts were only extracted where the marker used identified all transplanted cells (almost exclusively human nuclear markers) and not a specific subset of lineage restricted cells. For dopaminergic differentiation, cell counts extracted were based on tyrosine hydroxylase immunostaining or, in a limited number of studies, GFP expression driven by the dopaminergic *PITX3* promoter. In most articles, the cell numbers were provided directly by the authors, but in some articles, we estimated them from ranges and/or figures provided. Any estimated cell number is highlighted in Table 1 by <u>underlining</u>. Many articles employed transplant groups with various treatments, but for consistency and comparability between studies, only the unmanipulated/control cohorts are included here. Data extraction was completed independently by both authors, and cross-checked for accuracy.

**Table 1.**
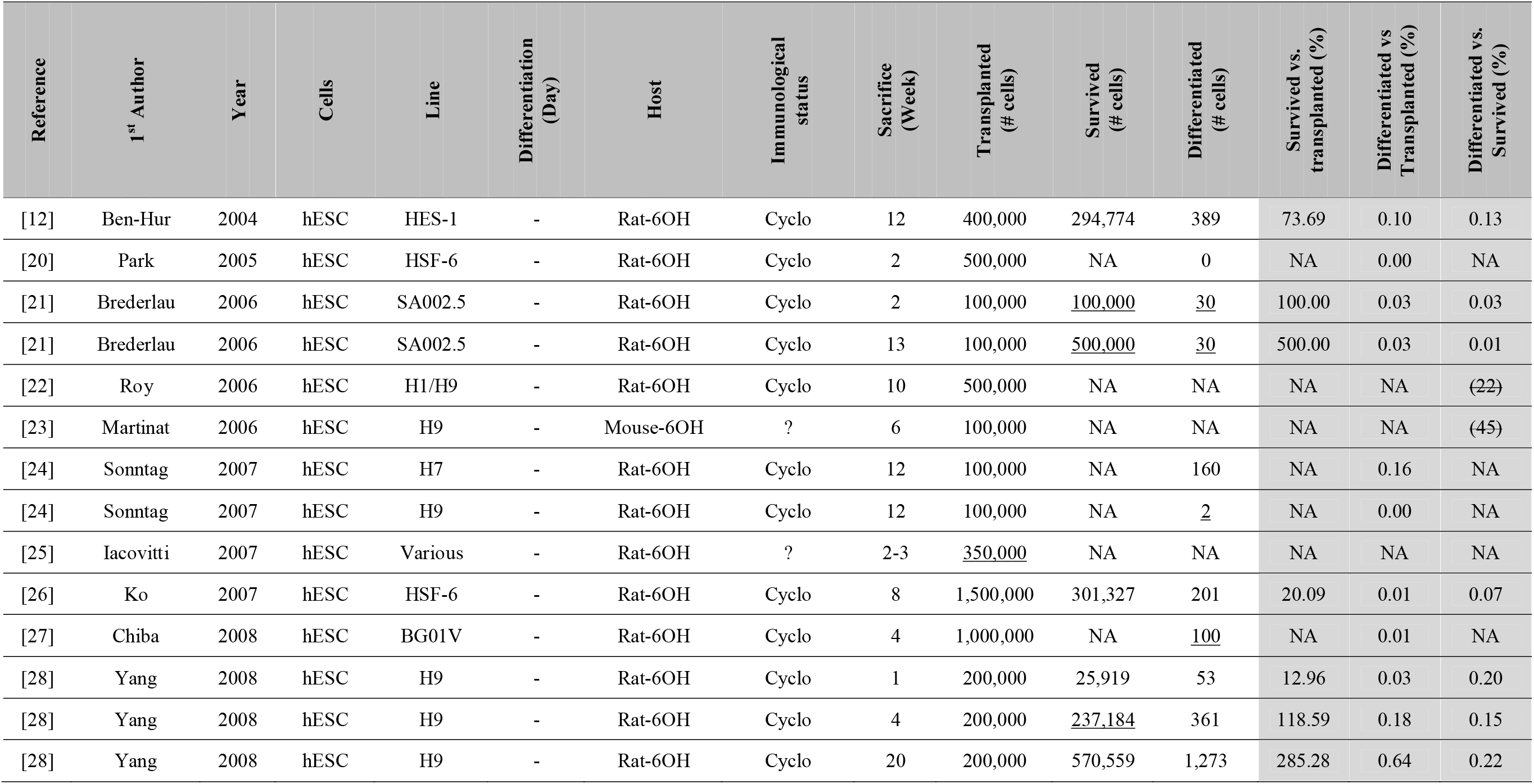
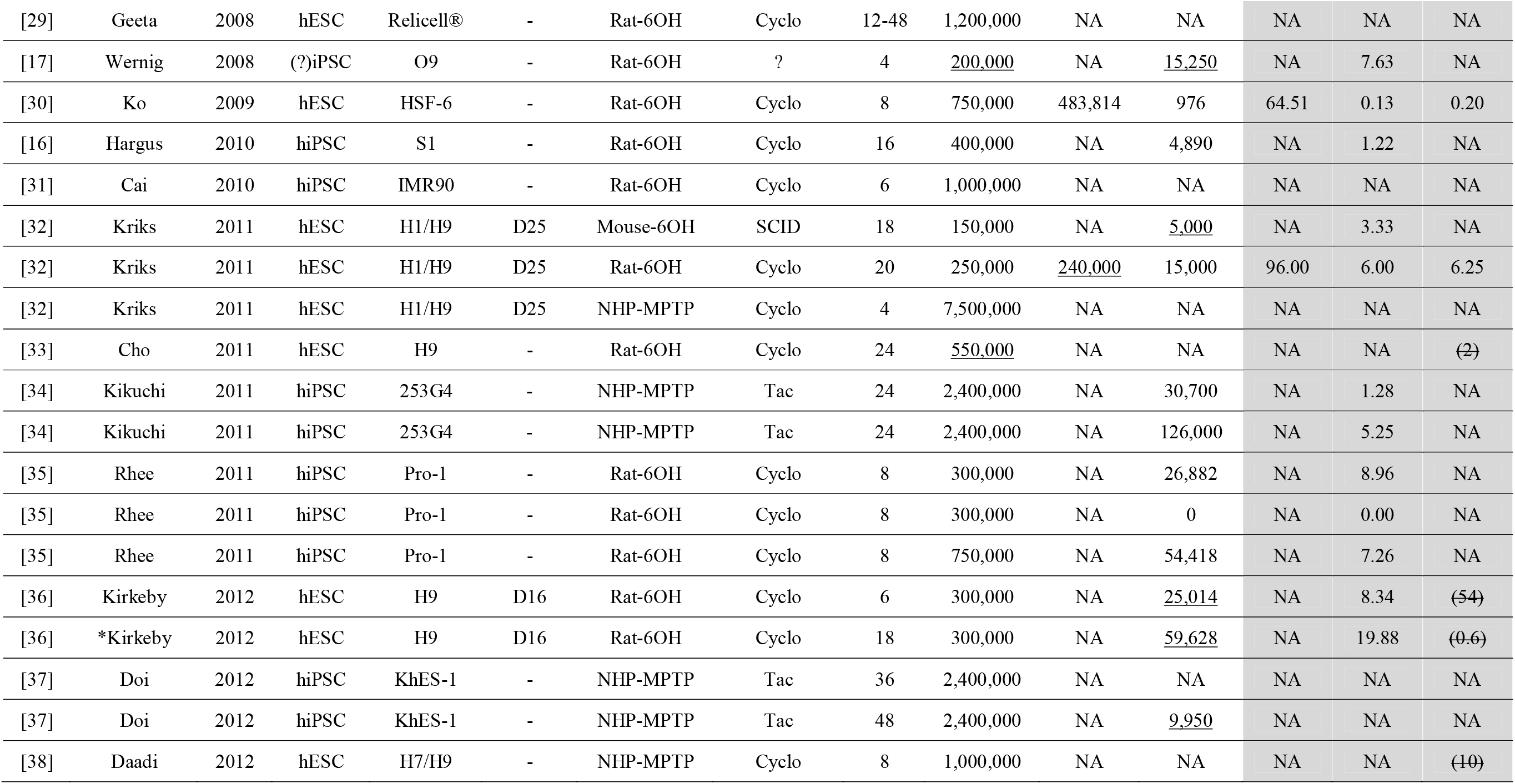
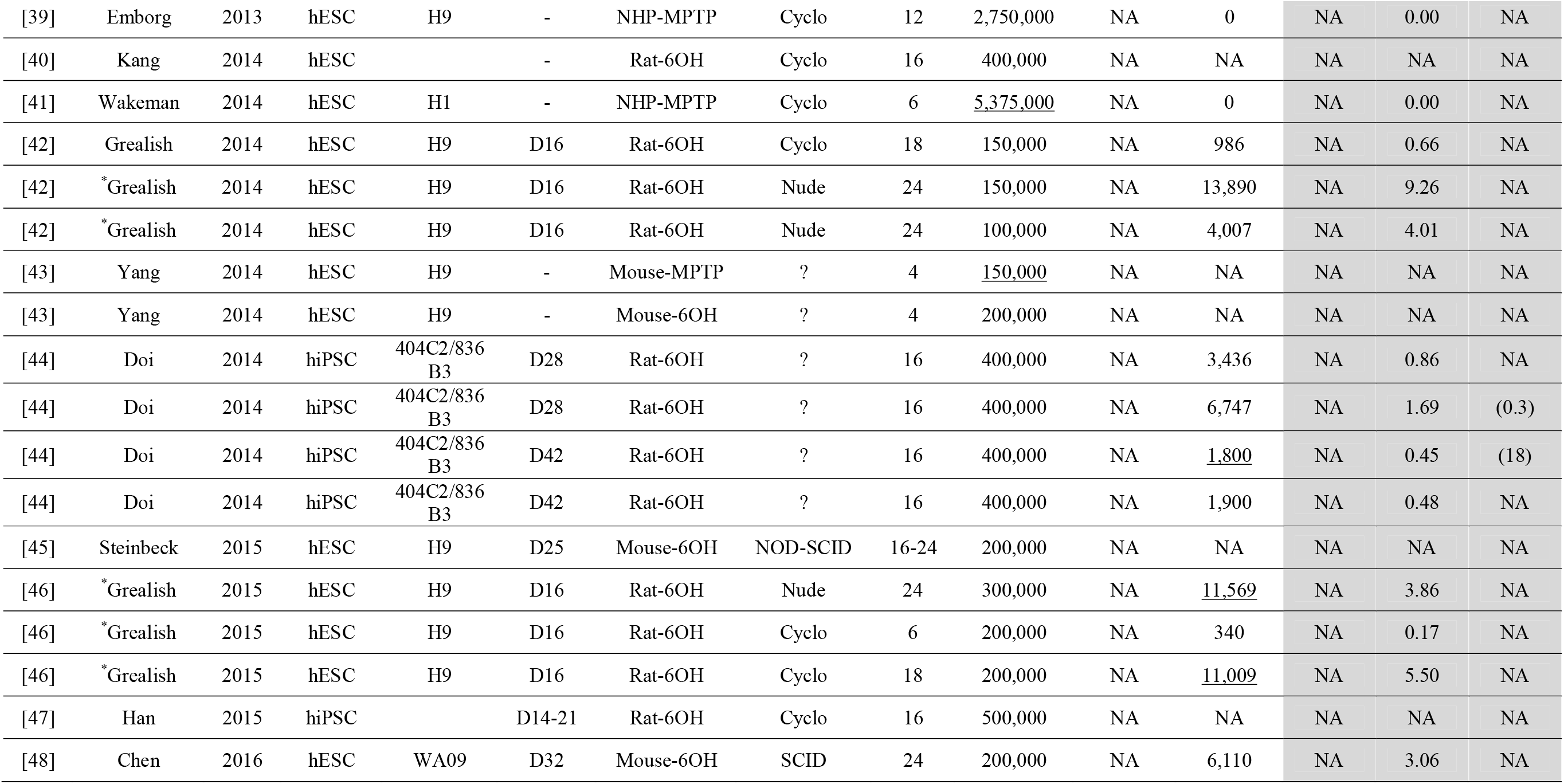
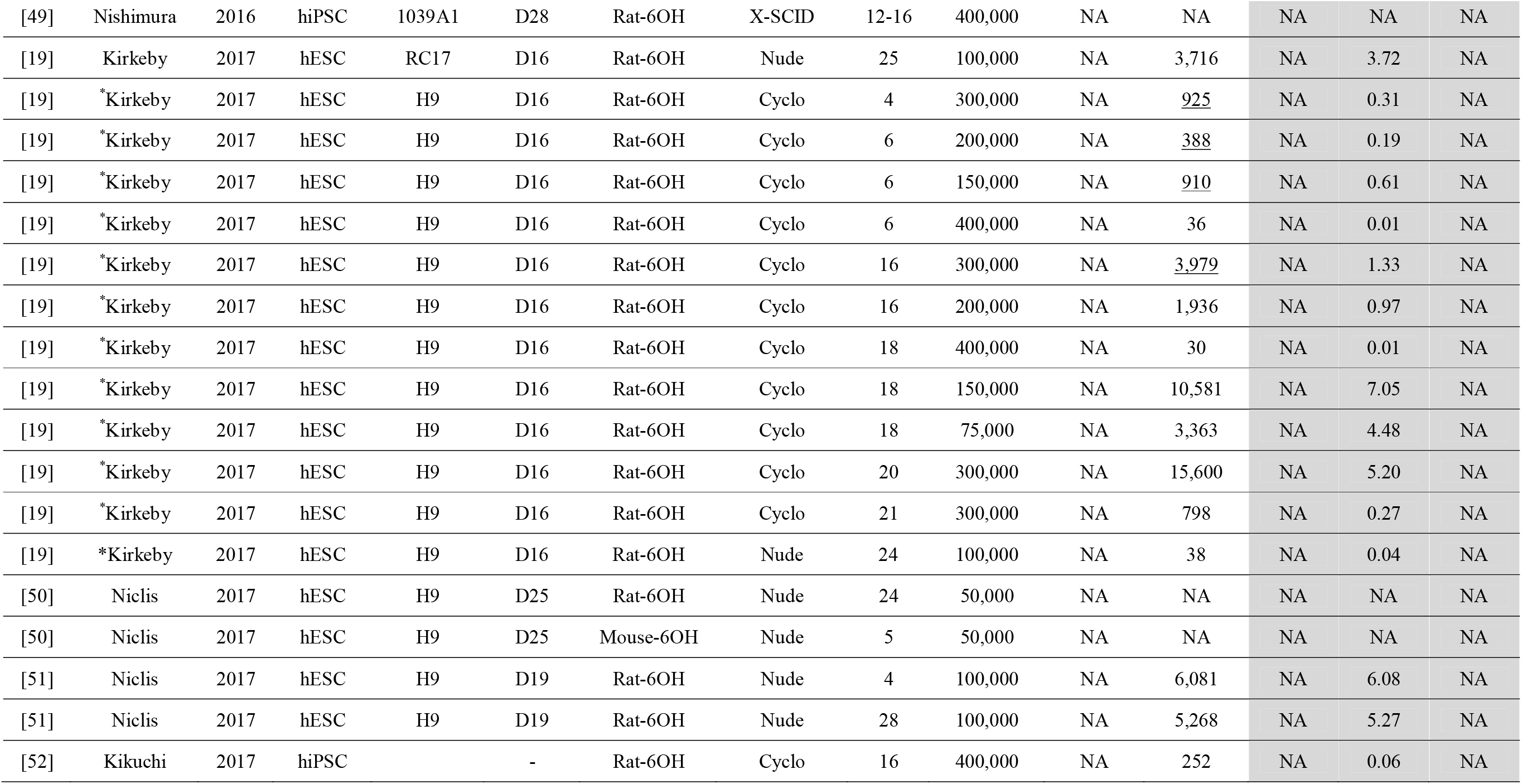
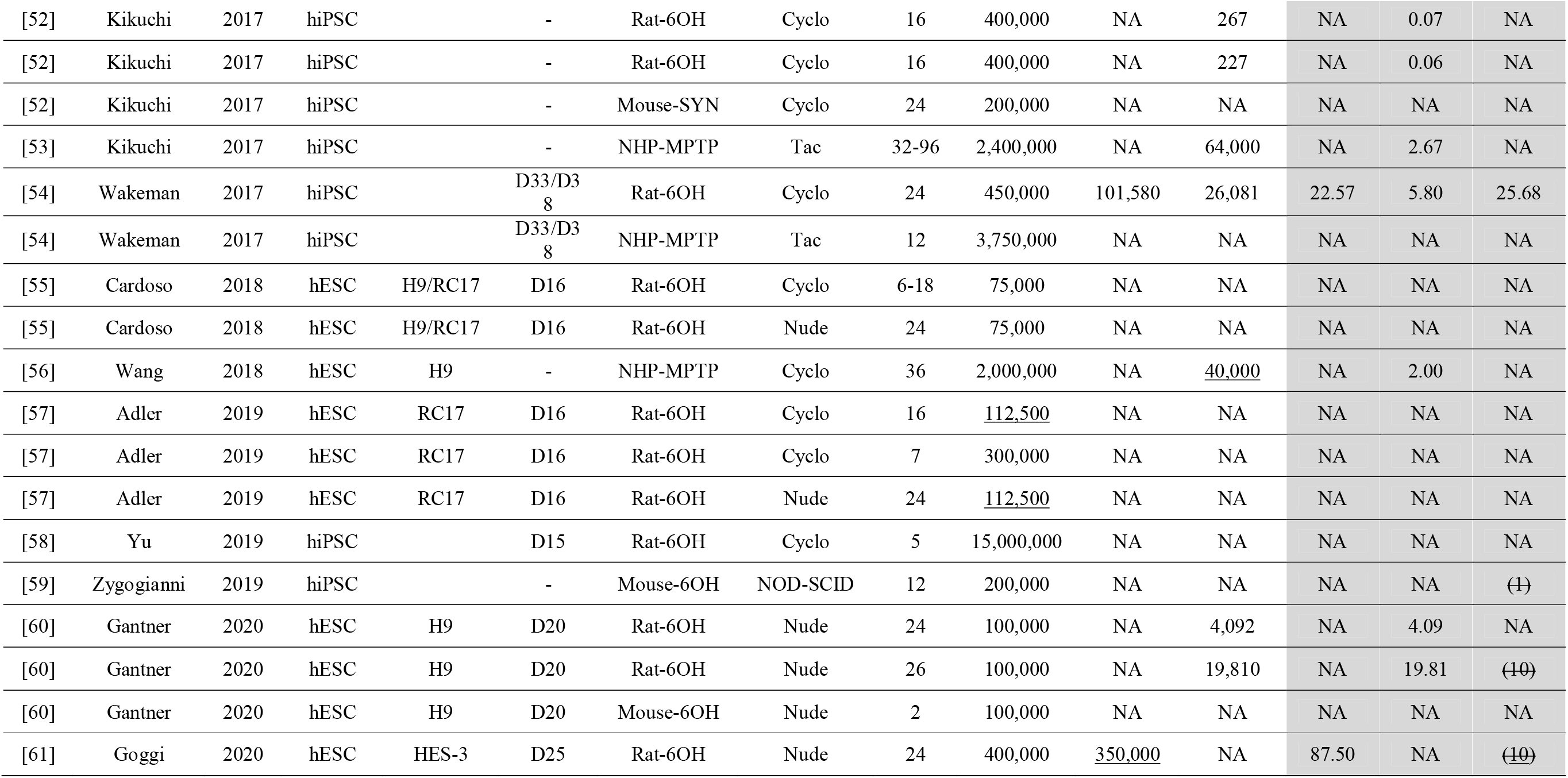
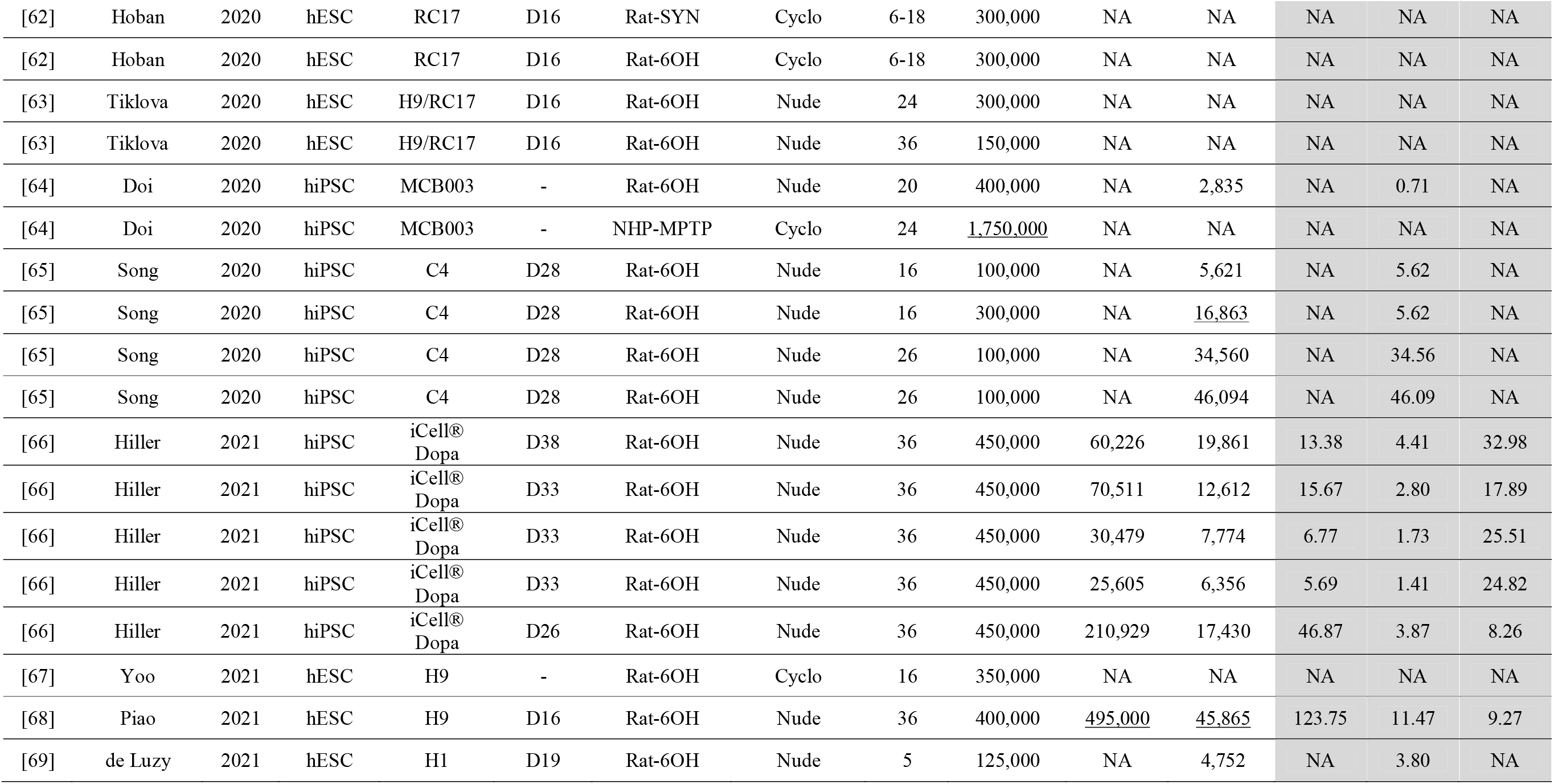
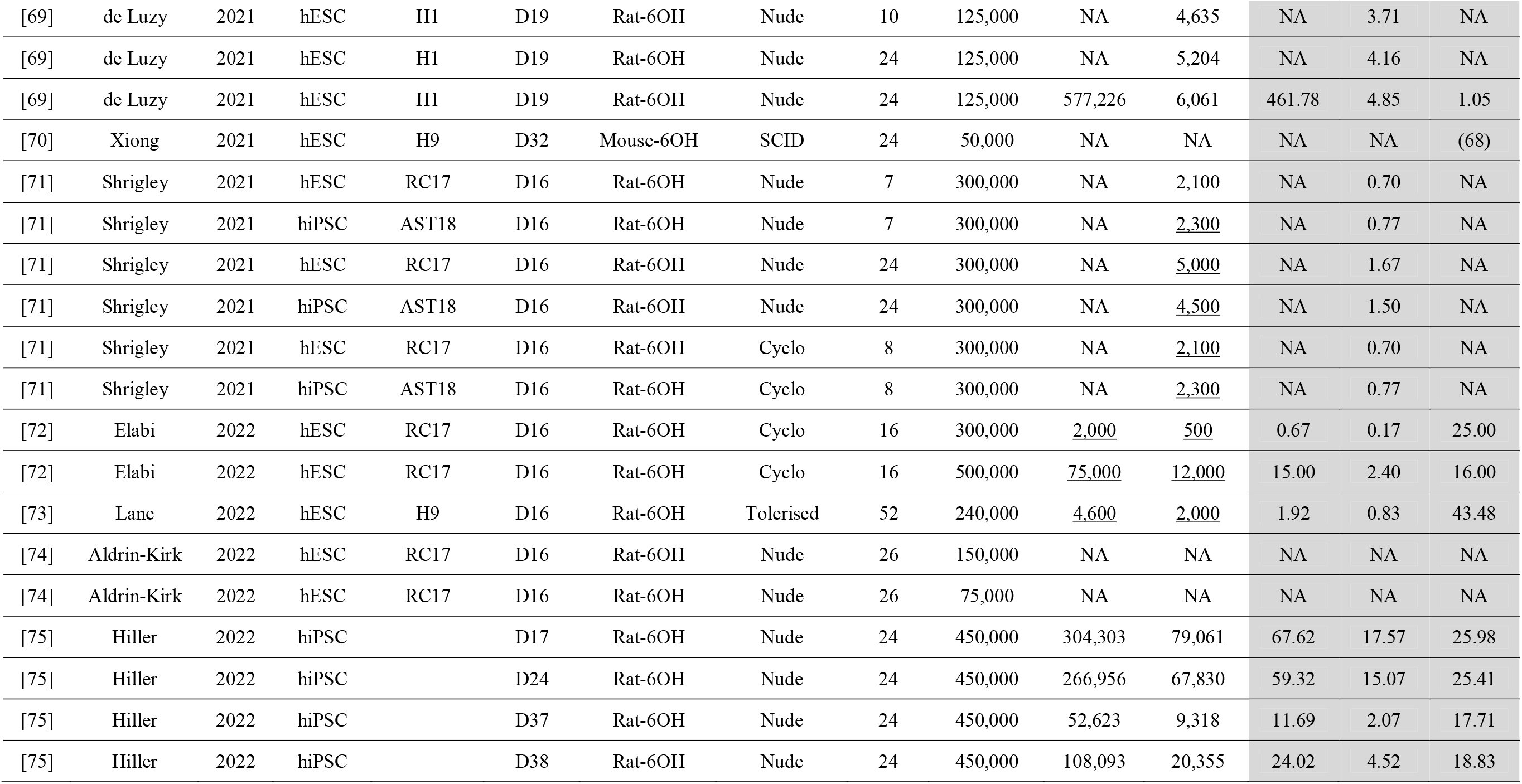
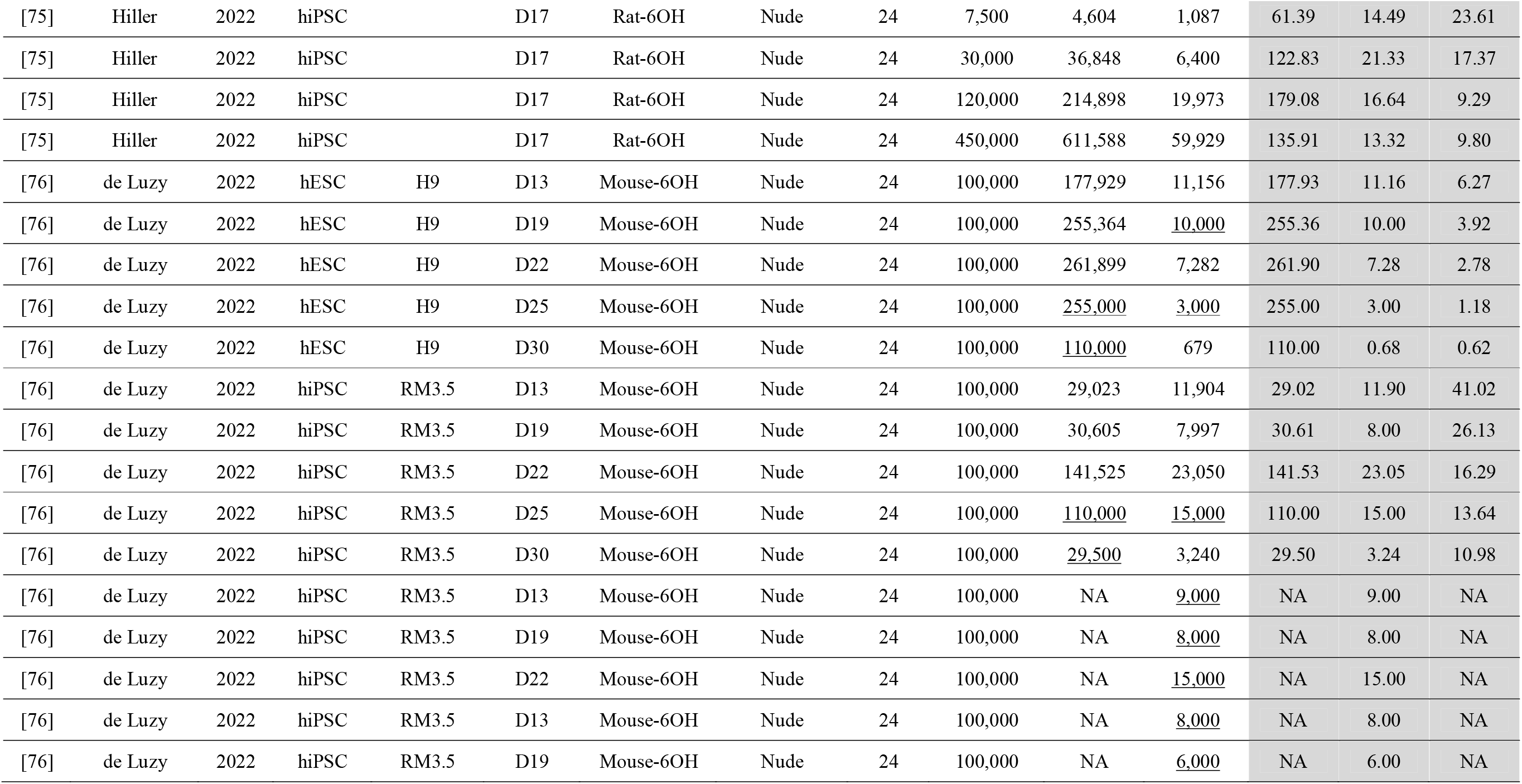
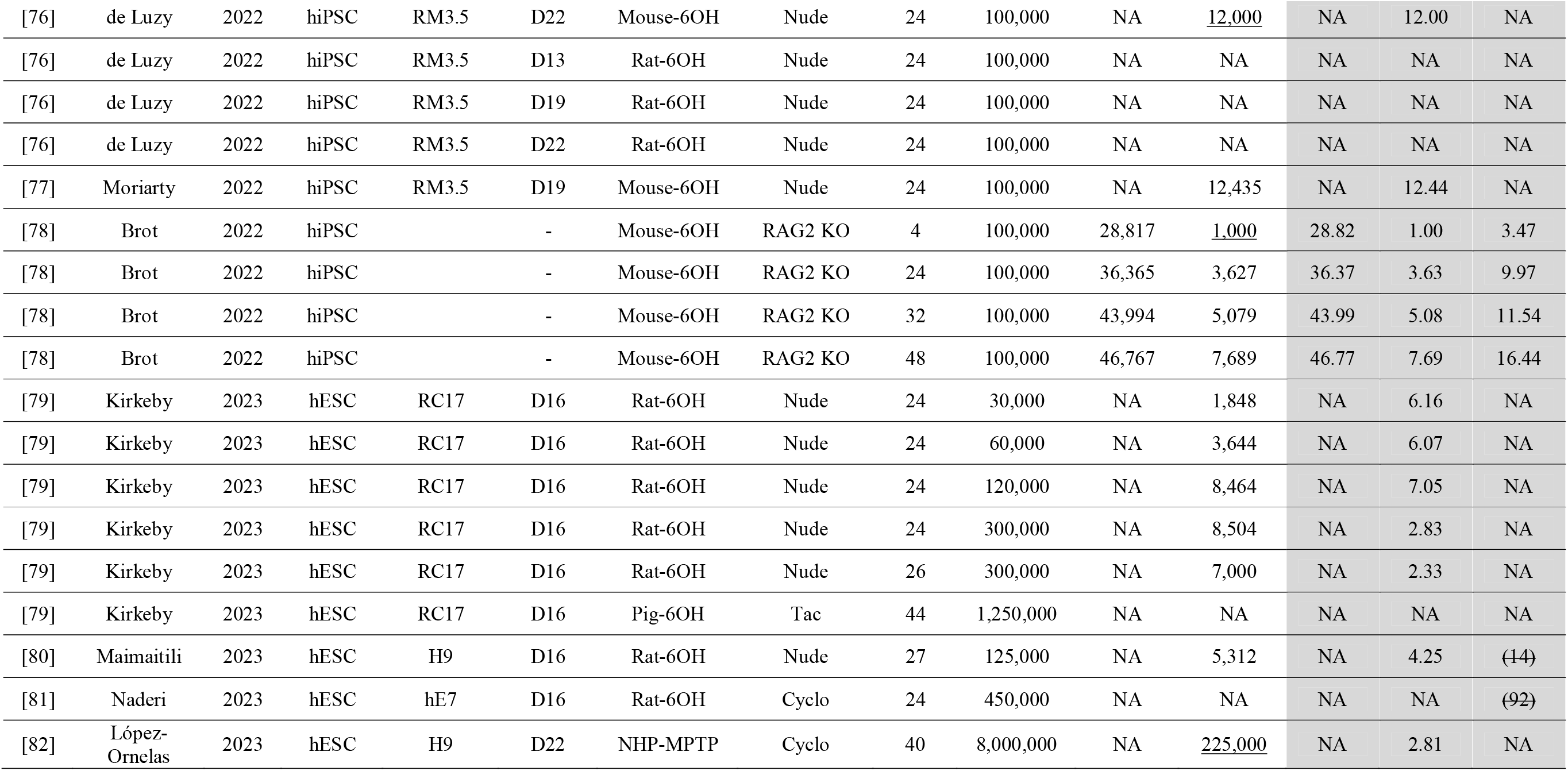
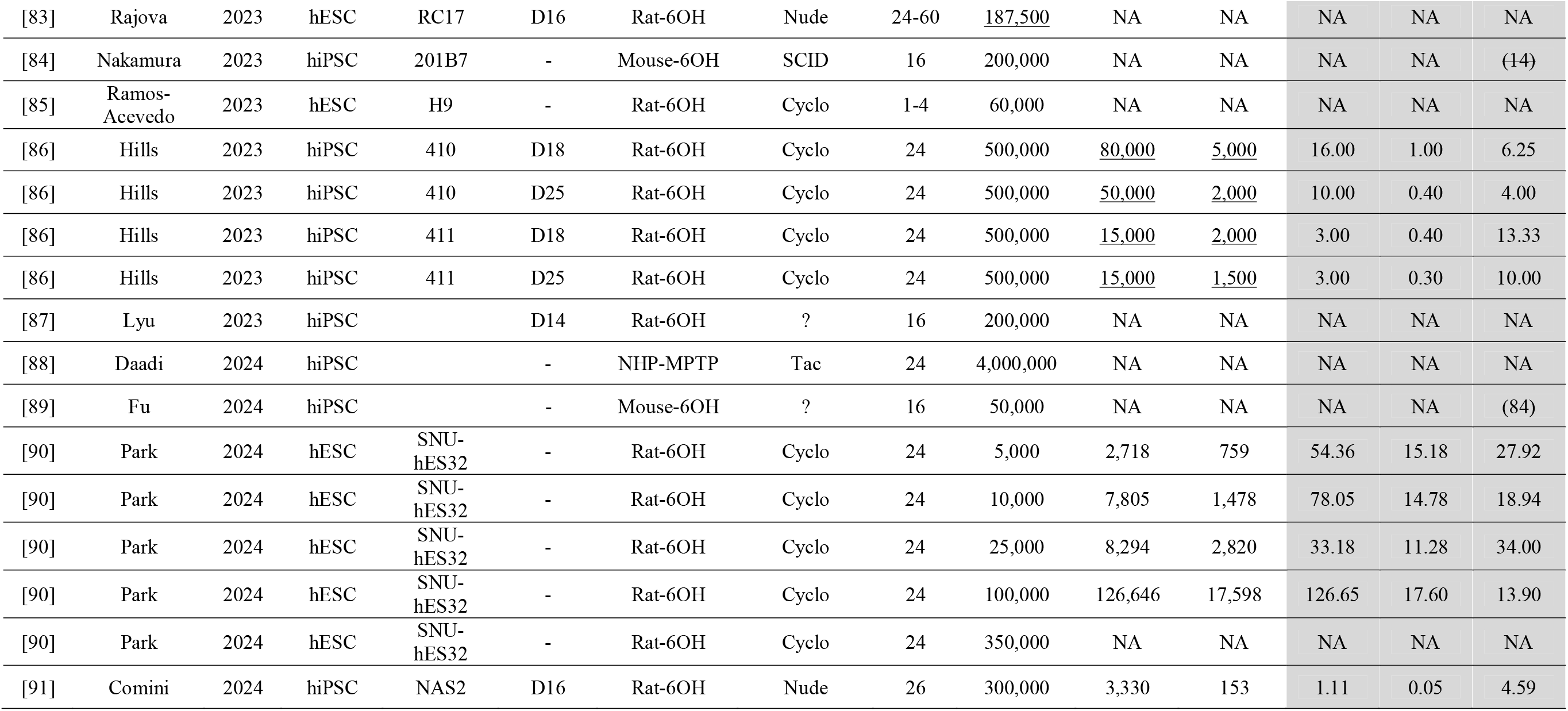
Studies included in this review. A total of 76 articles were included in this review from which 178 separate transplant studies were identified. The data extracted from these studies is depicted in this table. Any <u>underlined</u> value was estimated from a range or figure in the original article. Any number in brackets () was self-reported by the authors without any underpinning cell counts, and these were omitted from the results section. NA indicates that the data is not available for various reasons including not done, not reported, not a pan human cellular marker, expressed per section or per unit area/volume. *Indicates that the data was extracted from Table S1 in Kirkeby et al., 2017 [19]. Abbreviations: 6OH: 6-hydroxydopamine; SYN: Synuclein; Cyclo: Cyclosporine; Tac: Tacrine.

Based on the numbers of cells transplanted, surviving and differentiating, three parameters were then calculated: 1) cell survival (as a percent of cells transplanted, 2) cell differentiation (as a percent of cells transplanted) and 3) cell differentiation (as a percent of cells surviving). The latter was not depicted in any graphical representations in the results section but is available in Table 1. In a small number of articles, these percentages were self-reported by the authors without any underpinning cell counts. These are included (in brackets) in Table 1 but are not included in any graphical depictions of the data.

We also determined which articles included behavioural evaluation of graft-derived motor recovery (Supplementary Table 1) and, if so, which behavioural test was used and whether or not there was functional recovery.

### Limitations of this Review

This systematic review is intended to give an overview of survival and differentiation of human stem cell-derived dopaminergic progenitor transplantation in animal models of Parkinson’s. However, there are numerous factors that can affect graft survival and differentiation, and it is beyond the scope of the review to consider all of these. These factors include, but are not limited to, quality of stem cell line used, dopaminergic differentiation protocol used, validation of differentiation protocol, stage of dopaminergic differentiation at time of transplantation, verification of dopaminergic progenitor status at time of transplantation, use of cryopreservation for storage, viability of cell suspension used for transplantation, host animals used (species, strain, sex, age), immunological status of host animals (immunosuppressed or immunodeficient; immunosuppressive drug and drug regime; immunodeficient strain etc.), model of Parkinson’s induced, site of implantation, cell proliferation in the brain, time of sacrifice after transplantation, and the experience and expertise of the laboratories and individual researchers performing all of these tasks. Some, but not all, of these factors were recorded in Table 1. Another limitation is that many studies excluded animals from their analyses where there were no surviving grafts or no differentiated dopaminergic neurons. Therefore, the survival and differentiation data shown in the results section here may be an overrepresentation of the true figures.

### Statistical analysis of data

General study characteristics are shown as pie charts. All data related to cell survival and/or differentiation is shown as scatter plots depicting individual studies with the median. Any data cited in the text of the results section represents the mean ± standard error of the mean (SEM) unless specified otherwise. Comparisons between 2 groups were completed using Student’s *t*-test, comparisons between 3 or more groups on a single factor were completed using one-way ANOVA, and comparisons between groups on two factors were completed using two-way ANOVA. Where appropriate, *post-hoc* testing was applied, and the test used is specified in the corresponding figure. Simple linear regression was used to assess correlations in the data. Importantly, when considering the graphical depictions and the statistical analyses, it is important to bear in mind that there are many different variables (some of which are outlined above) that could underpin and affect the data shown.

## Results

### Extracted Data

From the 76 articles included in this review, 178 different transplant studies were extracted (Table 1).

### Study Characteristics

Of the 178 different transplant studies included in this review, 103 used human ESCs for transplantation while 75 used human iPSCs (Fig 2A). The vast majority of transplants were into 6-hydroxydopamine-lesioned rodents (128 rat studies and 32 mouse studies), with an additional 14 studies transplanting into MPTP-lesioned non-human primates and 4 into other models (1 rat α-synuclein model, 1 mouse α-synuclein model, 1 mouse MPTP model and 1 minipig 6-hydroxydopamine model) (Fig. 2B). An approximately equal number of studies used immunosuppressed host animals (81 studies) and immunodeficient host animals (85 studies), while 1 study used immune-tolerised hosts and the remainder (11 studies) did not specify (Fig. 2C).

**Fig. 2.**
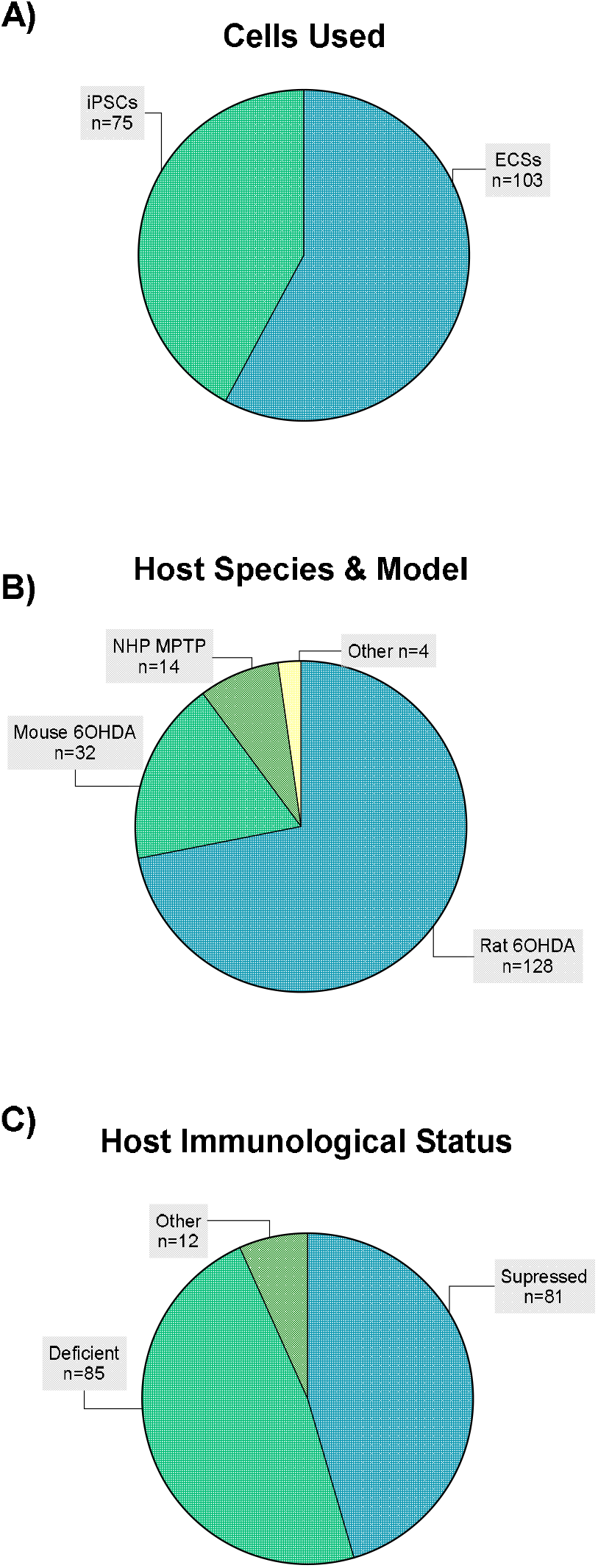
Study characteristics. Within the 76 articles included, 178 separate transplant studies were identified. The pie charts above show the proportions of these studies using A) different stem cells, B) host species and Parkinsonian models, and 3) the immunological status of the hosts.

### Studies reporting transplanted cell survival and/or differentiation

Of the 178 transplant studies, the number of cells surviving the transplant procedure was directly reported (or calculable from the ranges/figures provided) in 52 studies, while the number of cells differentiating into dopaminergic neurons was directly reported (or calculable) in 129 studies (Fig. 3). Since the number of cells transplanted was reported (or calculable) in all studies, this meant that the cell survival (as a percent of cells transplanted) could be calculated in 52 studies, cell differentiation (as a percent of cells transplanted) could be calculated in 129 studies.

**Fig. 3.**
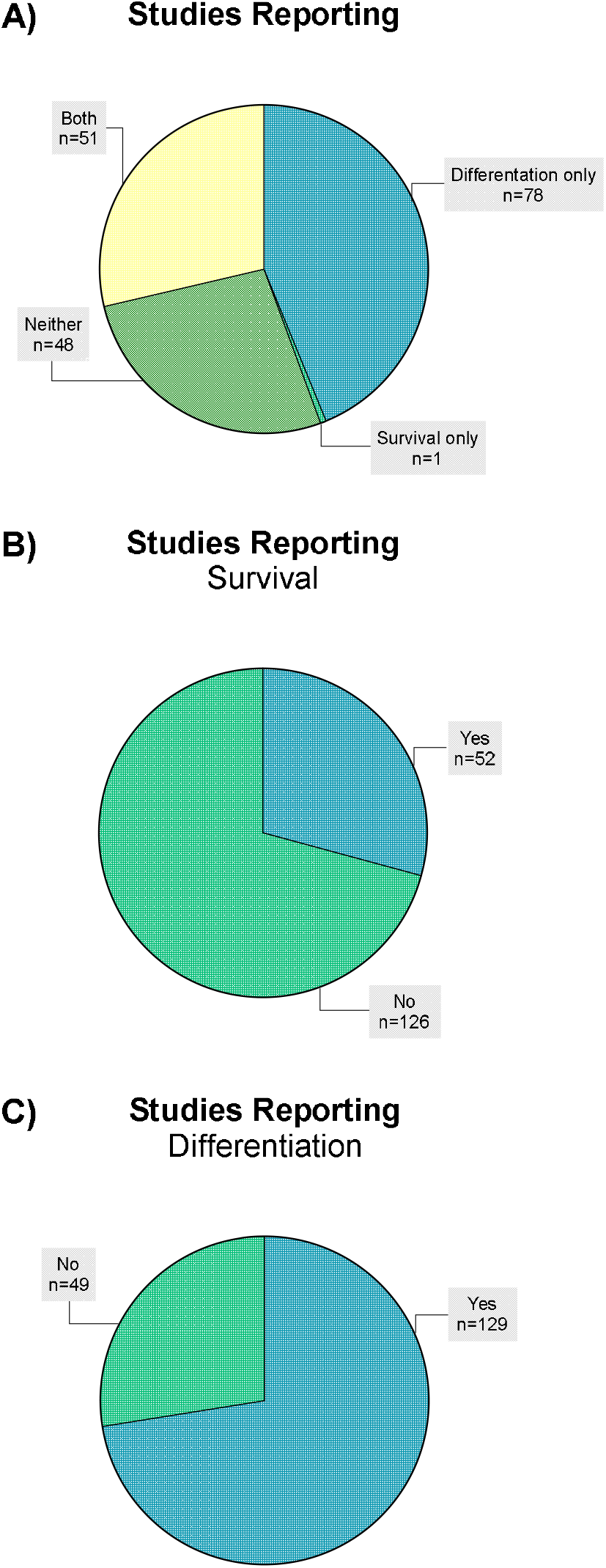
Studies reporting transplanted cell survival and/or differentiation. Within the 76 articles included, 178 separate transplant studies were identified. The pie charts above show the proportions of these studies reporting A & B) transplanted cell survival and A & C) transplanted cell dopaminergic differentiation.

### Overview of cell survival and differentiation

The overall survival of the cells transplanted in shown in Fig. 4A. This ranged from <1% to 500% of cells transplanted, with a median of 51% of transplanted cells surviving in the brain. Of note, 16 studies reported more cells surviving than were transplanted (i.e. >100%) indicating some level of cell proliferation after transplantation. The overall dopaminergic differentiation of the cells ranged from 0% to 46% of cells transplanted with a median of 3% (Fig. 4B). Interestingly, the dopaminergic differentiation was higher in proliferative grafts (with survival >100%) rather than non-proliferative grafts (with survival <100%) (Fig. 4C: Survival unknown: 4.7±0.8%; Survival <100%: 4.8±0.9%; Survival >100%: 9.8±1.9%; Cohort, *F*_(2,126)_ = 3.76, *P*<0.05).

**Fig. 4.**
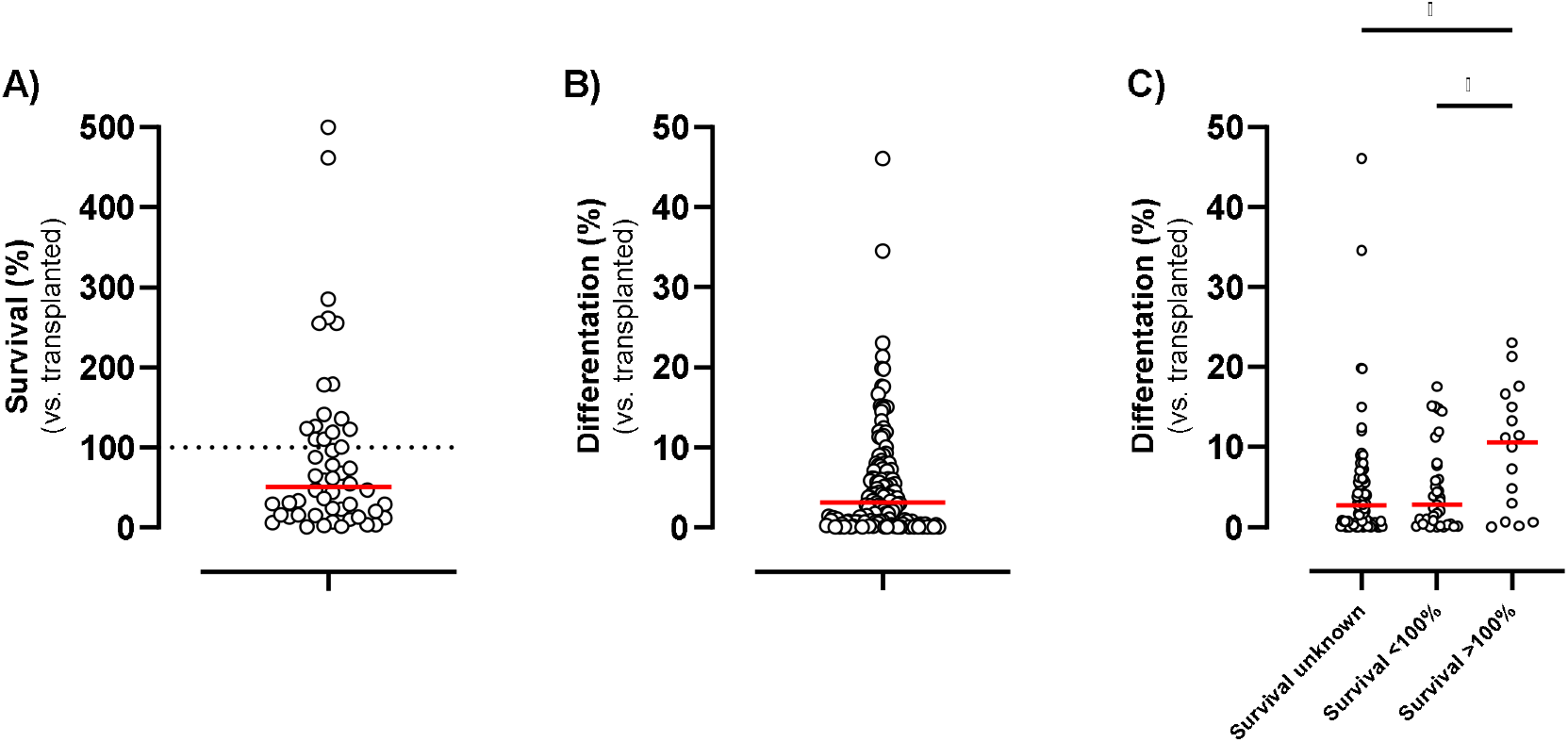
Overview of cell survival and differentiation. Of the 178 transplant studies extracted, 52 studies reported the number of cells surviving and 129 studies reported the number of dopaminergic neurons in the grafts. Thus, the scatter plots represent A) cell survival (as a percent of cells transplanted), B) dopaminergic differentiation (as a percent of cells transplanted) and C) dopaminergic differentiation (by survival cohort). Each point represents a separate transplant study, and the line indicates the median. Data were analysed by one-way ANOVA (C) with *post-hoc* Tukey’s and \**P*<0.05.

### Impact of cell type, host species and host immunological status on graft differentiation

Since *in situ* dopaminergic differentiation of transplanted stem cell-derived progenitors is a critical factor for effective brain repair, we next investigated whether some of the main experimental parameters affected this outcome. With regard to stem cell types used, there was a small but significantly higher dopaminergic differentiation in grafts derived from iPSCs compared with those derived from ESCs (Fig. 5A: ESCs: 4.1±0.6%; iPSCs: 6.9±1.1%; Cell type, *t*_(127)_ = 2.33, *P*<0.05). With regard to the host species, grafts in mice tended to have greater dopaminergic differentiation than those in rats or non-human primates (Fig. 5B: Rat: 5.0±0.8%; Mouse: 8.2±1.1%; NHP: 1.4±0.4%; Species, *F*_(2,126)_ = 3.33, *P*<0.05). However, this was only statistically significant on *post-hoc* analysis between mice and primates (*P*<0.05). In terms of the immunological status of the host animals, grafts in immunodeficient animals were significantly larger than those in immunosuppressed animals (Fig. 5C: Supressed: 2.8±0.6%; Deficient: 7.8±1.0%; Immunological status, *t*_(121)_ = 4.12, *P*<0.0001).

**Fig. 5.**
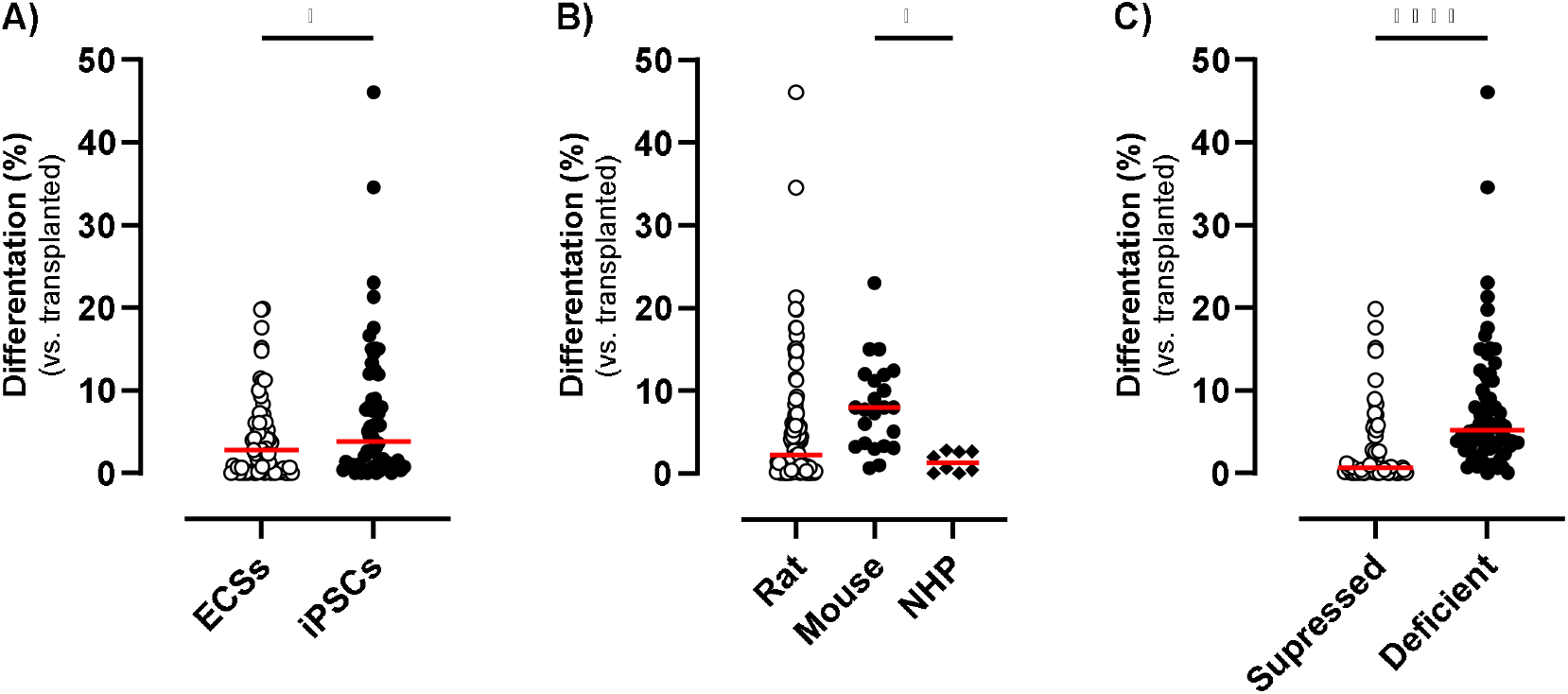
Impact of cell type, host species and host immunological status on graft differentiation. Of the 178 transplant studies extracted, 129 studies reported the number of dopaminergic neurons in the grafts. Thus, the scatter plots represent dopaminergic differentiation) by A) cell type, B) host species and C) host immunological status. Each point represents a separate transplant study, and the line indicates the median. Data were analysed by *t*-test (A and C) or one-way ANOVA with *post-hoc* Tukey’s (B) and \**P*<0.05, \*\*\*\**P*<0.0001.

### Further stratification of differentiation by cell type, host species and host immunological status

Since the data above suggested that stem cell type, host species and immunological status could affect dopaminergic differentiation, we next sought to further stratify the data to investigate this more. Since dopaminergic differentiation in non-human primates was very consistent and all animals were immunosuppressed, these were not considered in this section of the analysis.

We first compared ESC and iPSC-derived dopaminergic differentiation in immunosuppressed and immunodeficient rats (Fig. 6A). We found that there was no significant effect of cell type on differentiation (Cell type, *F*_(1,88)_ = 1.23, *P*>0.05), but there was a significant effect of immunological status with greater differentiation in immunodeficient animals (Immunological status, *F*_(1,88)_ = 9.80, *P*<0.01) and a significant interaction between the two (Cell type x Immunological status, *F*_(1,88)_ = 4.04, *P*<0.05). Subsequent *post-hoc* analysis confirmed that dopaminergic differentiation was greatest in iPSC-derived transplants in immunodeficient rats. We were unable to do a similar comparison for mice as all studies where differentiation was reported used immunodeficient mice. Rather, we next compared ESC and iPSC-derived dopaminergic differentiation between immunodeficient rats and mice (Fig. 6B). Interestingly, when all rodents were immunodeficient, we found that there was no significant effect of cell type or species on differentiation, and no interaction between the two.

**Fig. 6.**
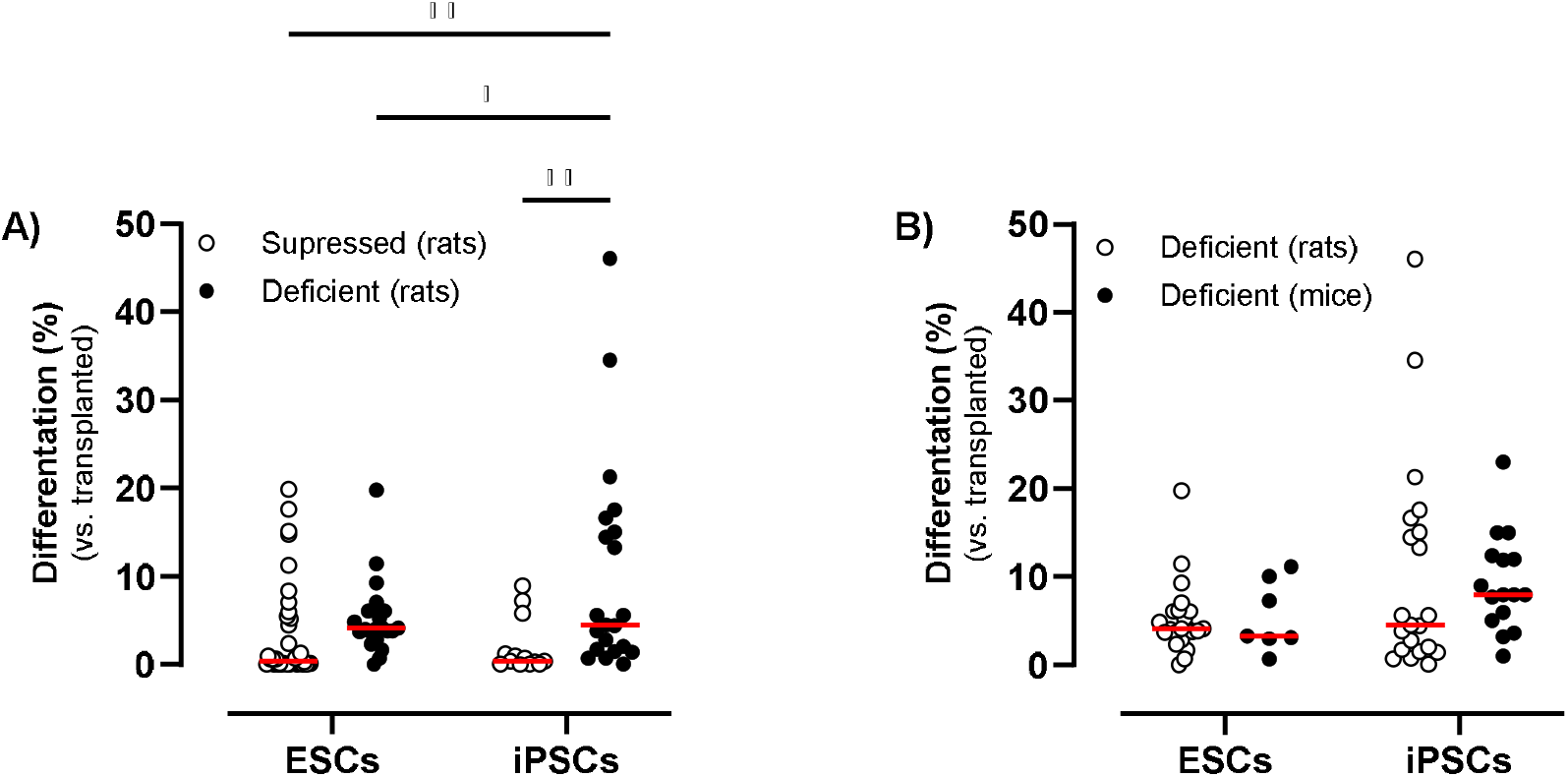
Further stratification of differentiation data in rodents. Of the 178 transplant studies extracted, 121 studies reported the number of dopaminergic neurons in the grafts in rats or mice. Thus, the scatter plots represent dopaminergic differentiation by A) cell type and immunological status in rats, and B) cell type in immunodeficient rats and mice. Each point represents a separate transplant study, and the line indicates the median. Data were analysed two-way ANOVA with *post-hoc* Newman Keuls and \**P*<0.05, \*\**P*<0.01.

### Assessment of graft functionality through behavioural testing

The most clinically relevant outcome of effective engraftment in Parkinsonian animals is amelioration of motor dysfunction. Therefore, we evaluated whether or not behavioural testing was included in the selected articles and studies therein (Supplementary Table 1). Of the 178 different transplant studies, 114 included at least one motor function test, while 64 did not do any behavioural testing (Fig. 7A). The most commonly-used behavioural test was amphetamine-induced rotation (used in 95 studies) followed by the cylinder test (used in 22 studies). In the studies using amphetamine-induced rotation, recovery was reported in 81% (Fig. 7B) and, not surprisingly, these had a significantly greater number of cells differentiating compared with the 19% of studies where amelioration of rotational asymmetry did not occur (Fig. 7C; Recovery, *F*_(1,79)_ = 2.35, *P*<0.05).

**Fig. 7.**
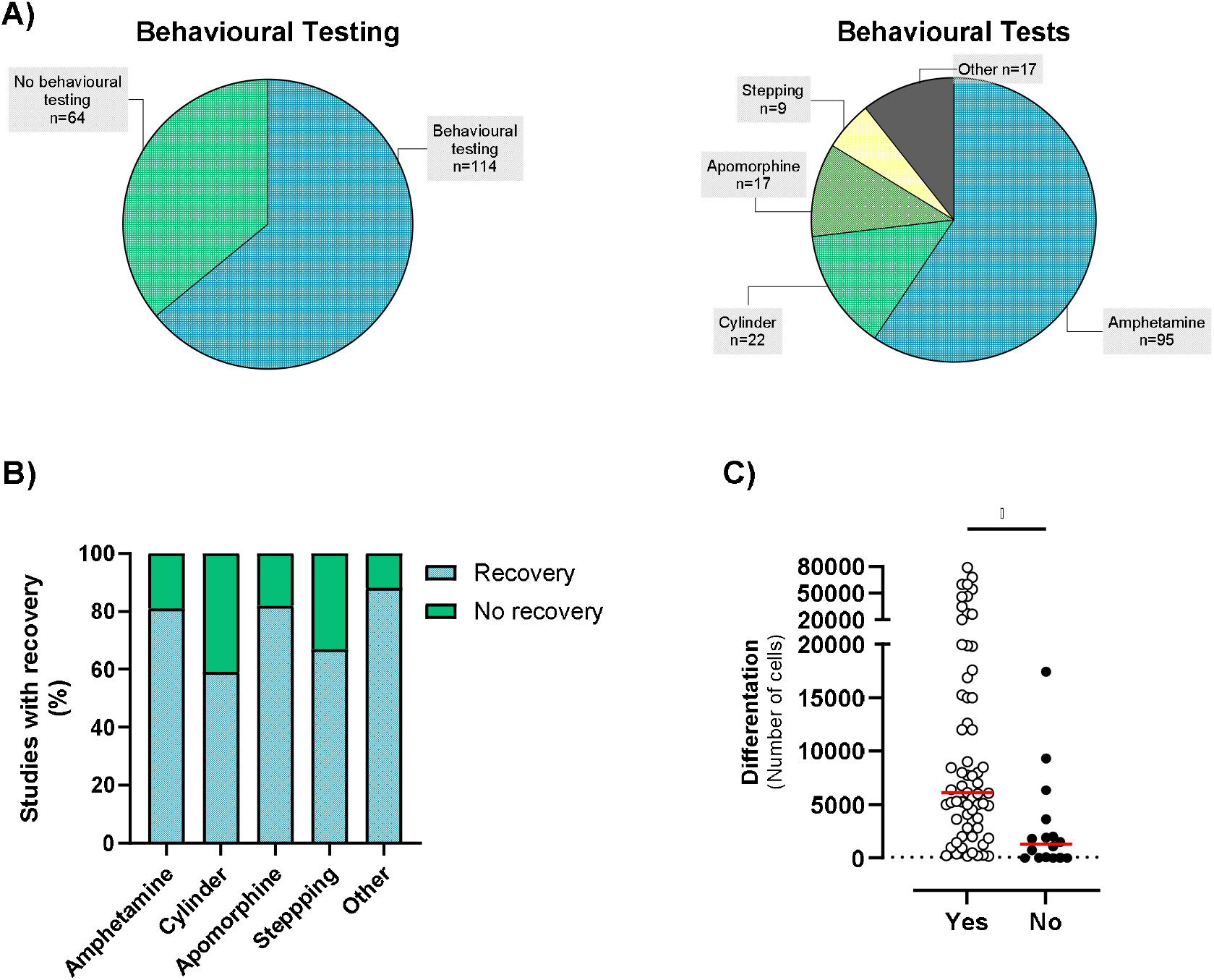
Assessment of graft functionality through behavioural testing. Of the 178 transplant studies extracted, 114 studies included behavioural testing (A). The most widely-used test was amphetamine-induced rotation with recovery in 81% of studies (B). The scatter plot represents dopaminergic differentiation in studies where amelioration of rotational asymmetry did, or did not, occur. Note that the lowest number of cells in the Yes column was 160. Each point represents a separate transplant study, and the line indicates the median. Data were analysed by *t*-test (C) and \**P*<0.05.

### Impact of other factors on graft survival and/or differentiation

To visualise some other trends in the data, we plotted graft survival and differentiation as a function of day of dopaminergic differentiation at the time of transplantation (for protocols using dual SMAD inhibition only; Fig. S1), week of sacrifice after transplantation (Fig. S2) and year of publication (Fig. S3). As expected, these show that most studies transplanted cells between day 16-19 of dopaminergic differentiation and sacrificed the animals around 6 months after transplantation. Interestingly, there was a significant positive correlation between year of publication and dopaminergic differentiation *in situ* in the brain (Fig. 3B; Regression, *F*_(1,127)_ = 14.50, R=0.32, *P*<0.001), suggesting that differentiation has improved overall in the two decades since the first publication. However, scrutiny of Table 1 indicates that this is driven by a high differentiation rate in just two articles each with several individual studies [75, 76]. When these were omitted from the analysis there was no correlation between year of publication and differentiation. We also plotted graft survival and differentiation as a function of the number of cells transplanted (for rodent studies only, Fig. S4). Interestingly, this showed a significant negative correlation between the number of cells transplanted and the final percentage of cells surviving (Fig. S4Aii; Regression, *F*_(1,49)_ = 8.37, R=0.38, *P*<0.01) and differentiating (Fig. S4Bii; Regression, *F*_(1,117)_ = 15.39, R=0.34, *P*<0.001), suggesting that larger transplants have an overall poorer engraftment outcome. Finally, we plotted graft differentiation as a function of graft survival and found a significant positive correlation (Fig. S5; Regression, *F*_(1,49)_ = 7.65, R=0.37, *P*<0.01), suggesting that if grafts survived well, they also tended to differentiate well.

## Discussion

Stem cell-derived brain repair for Parkinson’s offers the possibility of life-long reconstruction of the degenerated dopaminergic input to the striatum after a single surgical transplant [6]. Relative to fetal tissue transplants, stem cells can provide unlimited numbers of the required phenotype of dopaminergic neurons for allogenic transplantation, with the iPSCs offering the additional potential of autologous transplantation. The clinical success of ESC and/or iPSC-derived brain repair will ultimately depend on both the safety and efficacy of the approach, and the field is awaiting the outcome of the iPSC trial (Kyoto Trial: UMIN000033564) and the three ESC trials (BlueRock Trial: NCT04802733; STEM-PD: NCT05635409; S.Biomedics Trial: NCT05887466) in this regard. However, even though stem cell-derived brain repair has already entered clinical trial, we observed that the *in situ* survival and dopaminergic differentiation reported in preclinical studies was highly variable. Given the importance of these parameters in determining the efficacy of brain repair, we sought to complete this systematic review to shed light on the current status of these in human ESC or iPSC studies in Parkinsonian models.

With regard to survival of the stem cell-derived progenitors in the brain after transplantation, we were able to determine this in about a third of studies. In these studies, it was highly variable (ranging from <1% to 500% of cells transplanted) but relatively high overall (median of 51%). Several studies reported a greater number of cells surviving than were transplanted (reflective of *in situ* proliferation), and this was not confined to historical studies or to transplants of early-stage progenitors. Given the tumorigenicity risks posed by proliferative cells in grafts, it will be important to continue refining pre-transplant screening to reduce the number of such cells transplanted into the brain, but also to consider post-transplant failsafes such as the use of stem cell lines expressing suicide genes [69, 92].

The ability of transplanted dopaminergic progenitors to differentiate and mature into dopaminergic neurons after transplantation into the Parkinsonian brain underpins any functional recovery that will be afforded by brain repair. This was reported by a large proportion of the studies identified and was also found to be highly variable (ranging from 0% to 46% of cells transplanted), but unlike the high survival rate, differentiation was low overall (median of 3%). Interestingly, dopaminergic differentiation was higher in studies with proliferative grafts (survival >100%) which contrasts with suggestions that the dopaminergic yield may be lower in proliferative grafts due to an overabundance of unspecified cells [69]. Numerous factors can affect the post-transplant differentiation of dopaminergic progenitors (as outlined in the Methods section) and it was not possible to consider all of these parameters here. Nevertheless, we did stratify the differentiation data on some of these factors and found some interesting trends.

The initial stratification suggested that cell line, host species and host immunological status were all factors that affected dopaminergic transplantation, and subsequent analysis (in rodents only) suggested that transplanting progenitors into immunosuppressed animals negatively impaired their ability to differentiate into mature neurons compared with transplanting them into immunodeficient animals. If this were to translate to human studies, it might suggest that differentiation, and therefore clinical efficacy, may be impaired in donor-to-patient allogenic transplantation which requires the use of long-term immunosupression. Conversely, the higher differentiation rate in immunodeficient animals – the vast majority of which were athymic T-cell deficient nude rats or mice – might suggest that autologous transplantation, which will not initiate any T-cell response, may have a better clinical outcome.

Another interesting, and somewhat surprising, aspect of the data was the trend for both survival and dopaminergic differentiation to be negatively correlated with the number of cells transplanted. Overall, transplanting high numbers of cells led to a lesser proportion surviving and differentiating in the brain. This could reflect cellular competition for trophic support, micronutrients, oxygen and synaptic connections with consequent cell death leading to activation of the innate immune system which is known to impair graft differentiation [93]. In addition to all transplants being into immunosuppressed animals, this could also partly explain the relatively low differentiation in non-human primates all of whom had the largest numbers of cells transplanted (∼3.5M of which only 1.4% differentiated). Again, if one were to speculate on the translational relevance of this, it might suggest that micro-transplantation – with multiple deposits of smaller numbers of cells – might have a better outcome than large deposits.

Although four clinical trials are ongoing to test the safety, tolerability and efficacy of both ESC and iPSC-derived transplants, this systematic review highlights the variability in survival and differentiation in the published pre-clinical literature. It also suggests that inclusion of proliferative cells within the transplant is still an ongoing issue that could undermine graft safety, while the relatively low differentiation (and negative correlation between differentiation and cells transplanted) could undermine the efficacy of this approach. Overall, this systematic review indicates that there remains scope for improvement in the differentiation of stem cell-derived dopaminergic progenitors in order to maximize the therapeutic potential of this approach for patients.

## Acknowledgements

Our research in this field is supported by research grants from the Michael J Fox Foundation for Parkinson’s Research (Grant Numbers: 17244 and 023410) and Science Foundation Ireland (Grant Numbers: 19/FFP/6554).

