## Supplementary File for "Human stem cell transplantation for Parkinson’s disease: A systematic review of *in situ* survival and maturation of progenitors derived from human embryonic or induced stem cells in Parkinsonian models"

**Contents**

Supplementary Table 1: Studies with behavioural testing. Pages 2-5

Supplementary Fig. S1-S5. Pages 6-8

**Supplementary Table 1. Studies included in this review that incorporated behavioural testing.** Of the 178 different transplant studies, 99 included at least one motor function test. The information extracted from these studies is depicted in this table. Any underlined value was estimated from a range or figure in the original article. Abbreviations: 6OH: 6-hydroxydopamine; SYN: Synuclein. Rec: Recovery.

| **Reference** | **1st Author** | **Year** | **Host** | **Differentiated**  **(# cells)** | **Amphetamine** | **Cylinder** | **Apomorphine** | **Stepping** | **Other tests** |
| --- | --- | --- | --- | --- | --- | --- | --- | --- | --- |
| [12] | Ben-Hur | 2004 | Rat-6OH | 389 | Yes rec |  | Yes rec | Yes rec | Yes rec |
| [20] | Park | 2005 | Rat-6OH | 0 | No rec |  |  | No rec |  |
| [21] | Brederlau | 2006 | Rat-6OH | 30 | No rec |  |  |  |  |
| [21] | Brederlau | 2006 | Rat-6OH | 30 | No rec |  |  |  |  |
| [22] | Roy | 2006 | Rat-6OH | NA |  | No rec | Yes rec | Yes rec |  |
| [23] | Martinat | 2006 | Mouse-6OH | NA |  |  | Yes rec |  |  |
| [24] | Sonntag | 2007 | Rat-6OH | 160 | Yes rec |  |  |  |  |
| [24] | Sonntag | 2007 | Rat-6OH | 2 | No rec |  |  |  |  |
| [26] | Ko | 2007 | Rat-6OH | 201 | Yes rec |  |  |  |  |
| [27] | Chiba | 2008 | Rat-6OH | 100 | No rec |  | Yes rec |  |  |
| [28] | Yang | 2008 | Rat-6OH | 1,273 | Yes rec | No rec |  | No rec |  |
| [29] | Geeta | 2008 | Rat-6OH | NA |  |  | Yes rec | Yes rec |  |
| [17] | Wernig | 2008 | Rat-6OH | 15,250 | Yes rec |  |  |  |  |
| [30] | Ko | 2009 | Rat-6OH | 976 | Yes rec |  |  |  |  |
| [16] | Hargus | 2010 | Rat-6OH | 4,890 | Yes rec | No rec | Yes rec | No rec |  |
| [31] | Cai | 2010 | Rat-6OH | NA | No rec |  | No rec |  |  |
| [32] | Kriks | 2011 | Mouse-6OH | 5,000 | Yes rec |  |  |  |  |
| [32] | Kriks | 2011 | Rat-6OH | 15,000 | Yes rec | Yes rec |  | Yes rec |  |
| [33] | Cho | 2011 | Rat-6OH | NA |  | No rec | Yes rec |  |  |
| [34] | Kikuchi | 2011 | NHP-MPTP | 30,700 |  |  |  |  | Yes rec |
| [34] | Kikuchi | 2011 | NHP-MPTP | 126,000 |  |  |  |  | Yes rec |
| [35] | Rhee | 2011 | Rat-6OH | 26,882 | Yes rec |  |  |  |  |
| [35] | Rhee | 2011 | Rat-6OH | 0 | No rec |  |  |  |  |
| [35] | Rhee | 2011 | Rat-6OH | 54,418 | Yes rec |  |  |  |  |
| [36] | *Kirkeby | 2012 | Rat-6OH | 59,628 | Yes rec | Yes rec |  |  |  |
| [37] | Doi | 2012 | NHP-MPTP | NA |  |  |  |  | No rec |
| [37] | Doi | 2012 | NHP-MPTP | 9,950 |  |  |  |  | Yes rec |
| [40] | Kang | 2014 | Rat-6OH | NA |  |  | Yes rec |  |  |
| [42] | Grealish | 2014 | Rat-6OH | 986 | Yes rec |  |  |  |  |
| [43] | Yang | 2014 | Mouse-MPTP | NA |  |  |  |  | Yes rec |
| [43] | Yang | 2014 | Mouse-6OH | NA |  |  | Yes rec |  | Yes rec |
| [44] | Doi | 2014 | Rat-6OH | 3,436 | Yes rec |  |  |  |  |
| [44] | Doi | 2014 | Rat-6OH | 6,747 | Yes rec |  |  |  |  |
| [44] | Doi | 2014 | Rat-6OH | 1,800 | No rec |  |  |  |  |
| [44] | Doi | 2014 | Rat-6OH | 1,900 | No rec |  |  |  |  |
| [45] | Steinbeck | 2015 | Mouse-6OH | NA | Yes rec |  |  |  |  |
| [47] | Han | 2015 | Rat-6OH | NA |  |  | Yes rec |  | Yes rec |
| [48] | Chen | 2016 | Mouse-6OH | 6,110 | Yes rec | Yes rec |  |  | No rec |
| [49] | Nishimura | 2016 | Rat-6OH | NA | Yes rec |  |  |  |  |
| [19] | Kirkeby | 2017 | Rat-6OH | 3,716 | Yes rec | Yes rec |  |  |  |
| [50] | Niclis | 2017 | Rat-6OH | NA | Yes rec |  |  |  |  |
| [50] | Niclis | 2017 | Mouse-6OH | NA | Yes rec |  |  |  |  |
| [51] | Niclis | 2017 | Rat-6OH | 5,268 | Yes rec |  |  |  |  |
| [52] | Kikuchi | 2017 | Rat-6OH | 252 | Yes rec |  |  |  |  |
| [52] | Kikuchi | 2017 | Rat-6OH | 267 | Yes rec |  |  |  |  |
| [52] | Kikuchi | 2017 | Rat-6OH | 227 | Yes rec |  |  |  |  |
| [53] | Kikuchi | 2017 | NHP-MPTP | 64,000 |  |  |  |  | Yes rec |
| [54] | Wakeman | 2017 | Rat-6OH | 26,081 | Yes rec |  | Yes rec |  |  |
| [55] | Cardoso | 2018 | Rat-6OH | NA | Yes rec |  |  |  |  |
| [55] | Cardoso | 2018 | Rat-6OH | NA | Yes rec |  |  |  |  |
| [56] | Wang | 2018 | NHP-MPTP | 40,000 |  |  |  |  | Yes rec |
| [58] | Yu | 2019 | Rat-6OH | NA | No rec |  |  |  |  |
| [59] | Zygogianni | 2019 | Mouse-6OH | NA | Yes rec | Yes rec |  |  |  |
| [60] | Gantner | 2020 | Rat-6OH | 4,092 | Yes rec | No rec |  |  |  |
| [60] | Gantner | 2020 | Rat-6OH | 19,810 | Yes rec | No rec |  |  |  |
| [63] | Tiklova | 2020 | Rat-6OH | NA | Yes rec | Yes rec |  |  |  |
| [63] | Tiklova | 2020 | Rat-6OH | NA | Yes rec |  |  |  |  |
| [64] | Doi | 2020 | Rat-6OH | 2,835 | Yes rec |  |  |  |  |
| [65] | Song | 2020 | Rat-6OH | 5,621 | Yes rec |  |  |  |  |
| [65] | Song | 2020 | Rat-6OH | 16,863 | Yes rec |  |  |  |  |
| [65] | Song | 2020 | Rat-6OH | 34,560 | Yes rec | Yes rec |  | Yes rec | Yes rec |
| [65] | Song | 2020 | Rat-6OH | 46,094 | Yes rec | Yes rec |  | Yes rec | Yes rec |
| [66] | Hiller | 2021 | Rat-6OH | 19,861 | Yes rec |  |  |  |  |
| [66] | Hiller | 2021 | Rat-6OH | 12,612 | Yes rec |  |  |  |  |
| [66] | Hiller | 2021 | Rat-6OH | 7,774 | Yes rec |  |  |  |  |
| [66] | Hiller | 2021 | Rat-6OH | 6,356 | No rec |  |  |  |  |
| [66] | Hiller | 2021 | Rat-6OH | 17,430 | No rec |  |  |  |  |
| [67] | Yoo | 2021 | Rat-6OH | NA | Yes rec |  |  |  |  |
| [68] | Piao | 2021 | Rat-6OH | 45,865 | Yes rec |  |  |  |  |
| [69] | de Luzy | 2021 | Rat-6OH | 5,204 | Yes rec |  |  |  |  |
| [69] | de Luzy | 2021 | Rat-6OH | 6,061 | Yes rec |  |  |  |  |
| [70] | Xiong | 2021 | Mouse-6OH | NA | Yes rec | Yes rec |  |  | Yes rec |
| [71] | Shrigley | 2021 | Rat-6OH | 5,000 | Yes rec |  |  |  |  |
| [71] | Shrigley | 2021 | Rat-6OH | 4,500 | Yes rec |  |  |  |  |
| [72] | Elabi | 2022 | Rat-6OH | 500 | Yes rec |  | No rec |  |  |
| [72] | Elabi | 2022 | Rat-6OH | 12,000 | Yes rec |  | No rec |  |  |
| [73] | Lane | 2022 | Rat-6OH | 2,000 | Yes rec |  |  |  |  |
| [75] | Hiller | 2022 | Rat-6OH | 79,061 | Yes rec |  |  |  |  |
| [75] | Hiller | 2022 | Rat-6OH | 67,830 | Yes rec |  |  |  |  |
| [75] | Hiller | 2022 | Rat-6OH | 9,318 | No rec |  |  |  |  |
| [75] | Hiller | 2022 | Rat-6OH | 20,355 | Yes rec |  |  |  |  |
| [75] | Hiller | 2022 | Rat-6OH | 1,087 | No rec |  |  |  |  |
| [75] | Hiller | 2022 | Rat-6OH | 6,400 | Yes rec |  |  |  |  |
| [75] | Hiller | 2022 | Rat-6OH | 19,973 | Yes rec |  |  |  |  |
| [75] | Hiller | 2022 | Rat-6OH | 59,929 | Yes rec |  |  |  |  |
| [76] | de Luzy | 2022 | Mouse-6OH | 9,000 | Yes rec | Yes rec |  |  |  |
| [76] | de Luzy | 2022 | Mouse-6OH | 8,000 | Yes rec | Yes rec |  |  |  |
| [76] | de Luzy | 2022 | Mouse-6OH | 15,000 | Yes rec | No rec |  |  |  |
| [76] | de Luzy | 2022 | Mouse-6OH | 8,000 | Yes rec | Yes rec |  |  |  |
| [76] | de Luzy | 2022 | Mouse-6OH | 6,000 | Yes rec | Yes rec |  |  |  |
| [76] | de Luzy | 2022 | Mouse-6OH | 12,000 | Yes rec | No rec |  |  |  |
| [78] | Brot | 2022 | Mouse-6OH | 3,627 | No rec |  |  |  |  |
| [78] | Brot | 2022 | Mouse-6OH | 5,079 | Yes rec |  |  |  |  |
| [78] | Brot | 2022 | Mouse-6OH | 7,689 | Yes rec |  |  |  |  |
| [79] | Kirkeby | 2023 | Rat-6OH | 1,848 | Yes rec |  |  |  |  |
| [79] | Kirkeby | 2023 | Rat-6OH | 3,644 | Yes rec |  |  |  |  |
| [79] | Kirkeby | 2023 | Rat-6OH | 8,464 | Yes rec |  |  |  |  |
| [79] | Kirkeby | 2023 | Rat-6OH | 8,504 | Yes rec |  |  |  |  |
| [79] | Kirkeby | 2023 | Rat-6OH | 7,000 | Yes rec |  |  |  |  |
| [80] | Maimaitili | 2023 | Rat-6OH | 5,312 | Yes rec | No rec |  |  |  |
| [81] | Naderi | 2023 | Rat-6OH | NA | Yes rec |  |  |  |  |
| [82] | López-Ornelas | 2023 | NHP-MPTP | 225,000 |  |  |  |  | Yes rec |
| [84] | Nakamura | 2023 | Mouse-6OH | NA |  |  | Yes rec |  |  |
| [86] | Hills | 2023 | Rat-6OH | 5,000 | Yes rec |  |  |  |  |
| [86] | Hills | 2023 | Rat-6OH | 2,000 | No rec |  |  |  |  |
| [86] | Hills | 2023 | Rat-6OH | 2,000 | Yes rec |  |  |  |  |
| [86] | Hills | 2023 | Rat-6OH | 1,500 | No rec |  |  |  |  |
| [87] | Lyu | 2023 | Rat-6OH | NA |  |  | Yes rec |  |  |
| [88] | Daadi | 2024 | NHP-MPTP | NA |  |  |  |  | Yes rec |
| [89] | Fu | 2024 | Mouse-6OH | NA |  |  | Yes rec |  | Yes rec |
| [90] | Park | 2024 | Rat-6OH | 759 | No rec |  |  |  |  |
| [90] | Park | 2024 | Rat-6OH | 1,478 | Yes rec |  |  |  |  |
| [90] | Park | 2024 | Rat-6OH | 2,820 | Yes rec |  |  |  |  |
| [90] | Park | 2024 | Rat-6OH | 17,598 | Yes rec |  |  |  |  |

**Supplementary Figure S1-S4**

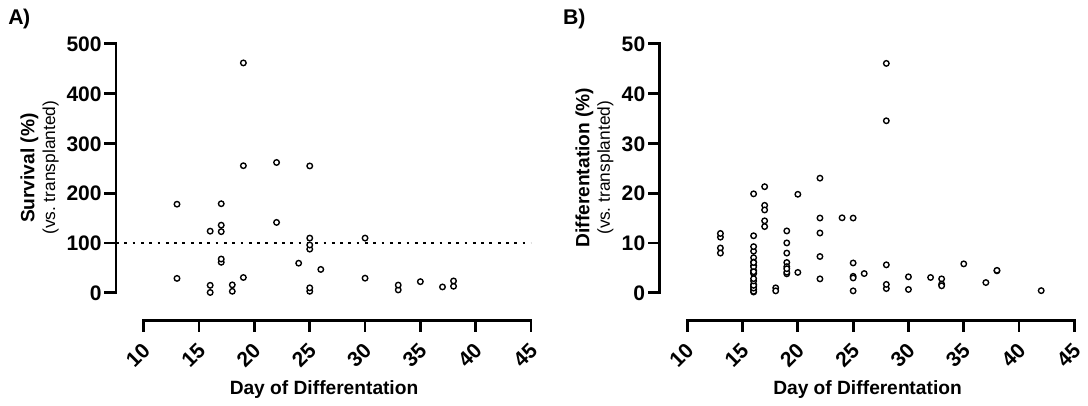

**Fig. S1 Effect of day of differentiation.** A) Graft survival and B) graft differentiation plotted as a function of day of dopaminergic differentiation at the time of transplantation (for protocols using dual SMAD inhibition only). This suggests that most studies transplanted cells between day 16-19 of dopaminergic differentiation.

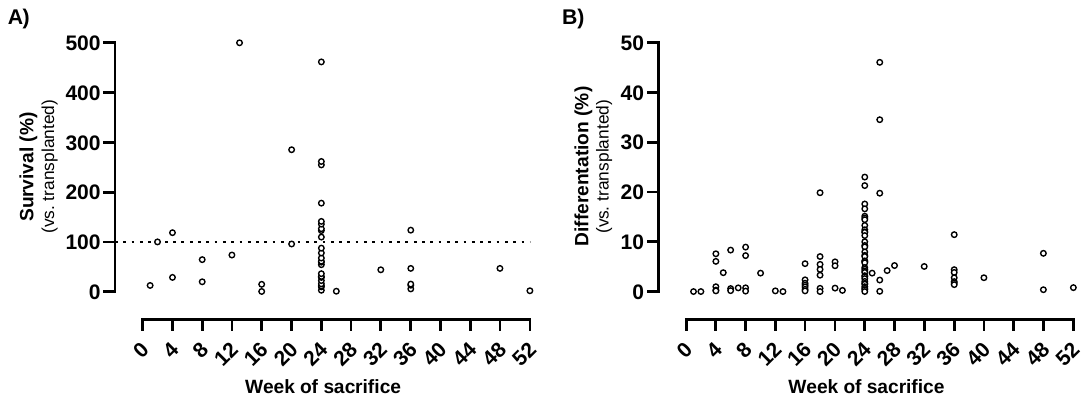

**Fig. S2 Effect of week of sacrifice.** A) Graft survival and B) graft differentiation plotted as a function of week of sacrifice after transplantation. This suggests that most studies sacrificed the host animals 6 months after transplantation.

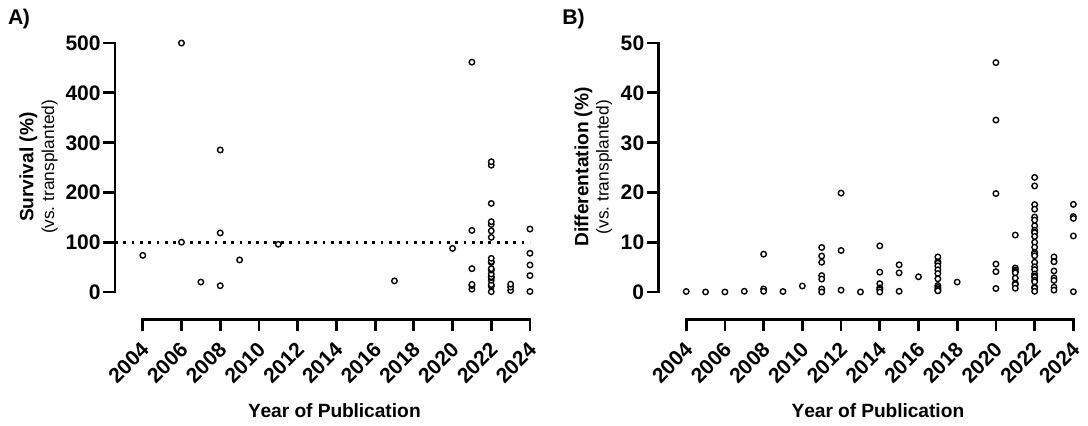

**Fig. S3 Effect of year of publication.** A) Graft survival and B) graft differentiation plotted as a function of year of publication. The increased differentiation in 2022 is driven by a high differentiation rate in just two articles (each with several individual studies).

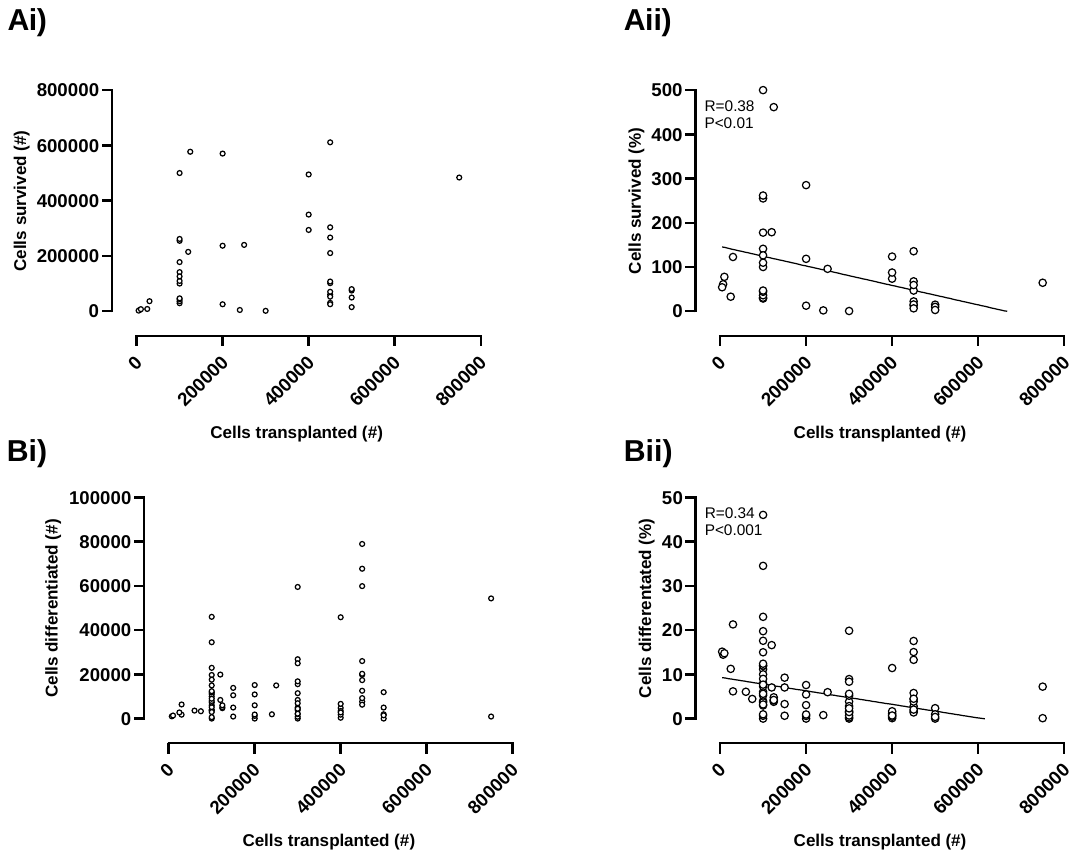

**Fig. S4 Effect of number of cells transplanted.** A) Graft survival and B) graft differentiation plotted as a function of cells transplanted. Note this data represents rodent grafts only and studies with >1M cells transplanted (n=2). are not shown because of x-axis scaling. Larger grafts have a poorer overall outcome.

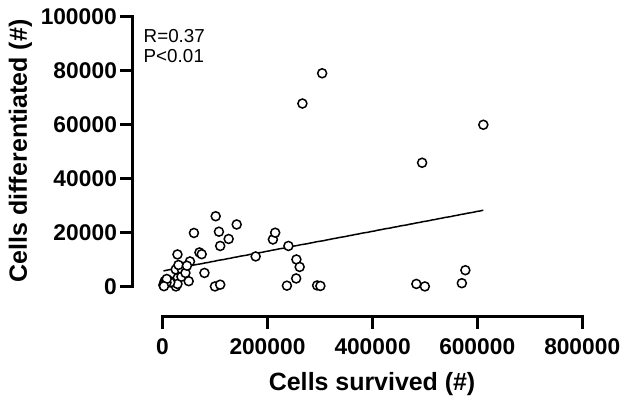

**Fig. S5 Effect of number of cells surviving.** Graft differentiation was plotted as a function of graft survival. This suggests that if grafts survived well, they also tended to differentiate well.
